# GelBox: Open-source software to improve rigor and reproducibility when analyzing gels and immunoblots

**DOI:** 10.1101/2024.03.07.583941

**Authors:** Utku Gulbulak, Austin G. Wellette-Hunsucker, Kenneth S. Campbell

## Abstract

GelBox is open-source software that was developed with the goal of enhancing rigor, reproducibility, and transparency when analyzing gels and immunoblots. It combines image adjustments (cropping, rotation, brightness, and contrast), background correction, and band-fitting in a single application. Users can also associate each lane in an image with metadata (for example, sample type). GelBox data files integrate the raw data, supplied metadata, image adjustments, and band-level analyses in a single file to improve traceability. GelBox has a user-friendly interface and was developed using MATLAB. The software, installation instructions, and tutorials, are available at https://campbell-muscle-lab.github.io/GelBox/.

## INTRODUCTION

Gel electrophoresis and immunoblotting are ubiquitous in many types of science. A PubMed search for [“gel electrophoresis” or “immunoblots”] returns 3,548 publications for 2023 alone. Many different experimental techniques can be used but they all produce an image that needs to be interpreted. A typical analysis involves cropping the image, adjusting the brightness and contrast, evaluating, and quantifying the bands. These operations can be performed using many different software tools.

Personal communications suggest that many scientists use workflows that involve one or more packages that include but are not restricted to Photoshop (Adobe, San Jose, CA), ImageQuant TL (GE Healthcare Bio-Sciences Corp., Marlborough, MA), Image Lab (Hercules, CA), ImageJ (1), Fiji (2), and IOCBIO Gel (3). Some scientists use one piece of software to crop and adjust an image and another program to analyze the bands. These diverse workflows provide flexibility but can be challenging to document and reproduce.

Our lab is committed to rigor and reproducibility and developed an in-house tool to help analyze our gels and blots. We wanted a system that integrated our typical workflows into a single package and allowed traceability. After discussing our approach with collaborators, we realized that our software might be helpful to other researchers. This paper details our approach and describes some of the strengths and limitations of the workflow.

## METHODS

GelBox (Figure 1) is a program developed in MATLAB App Designer (MathWorks, Natick, MA). The analysis workflow with GelBox starts with an image file and is illustrated in Figure 2.

**Figure 1.**
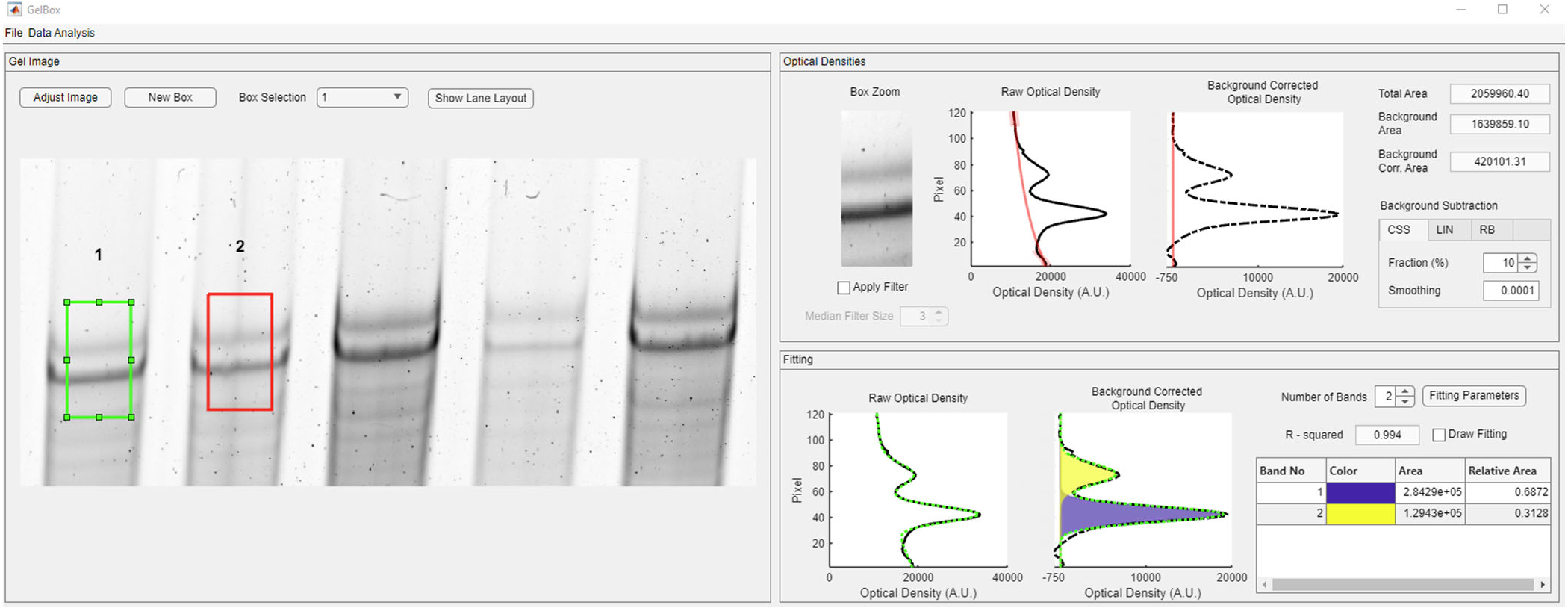
GelBox user interface. The green box shows the selected ROI and the resulting density profiles. The curve-fits are automatically calculated. The red box shows the second ROI on the gel. A.U. stands for arbitrary units.

**Figure 2.**
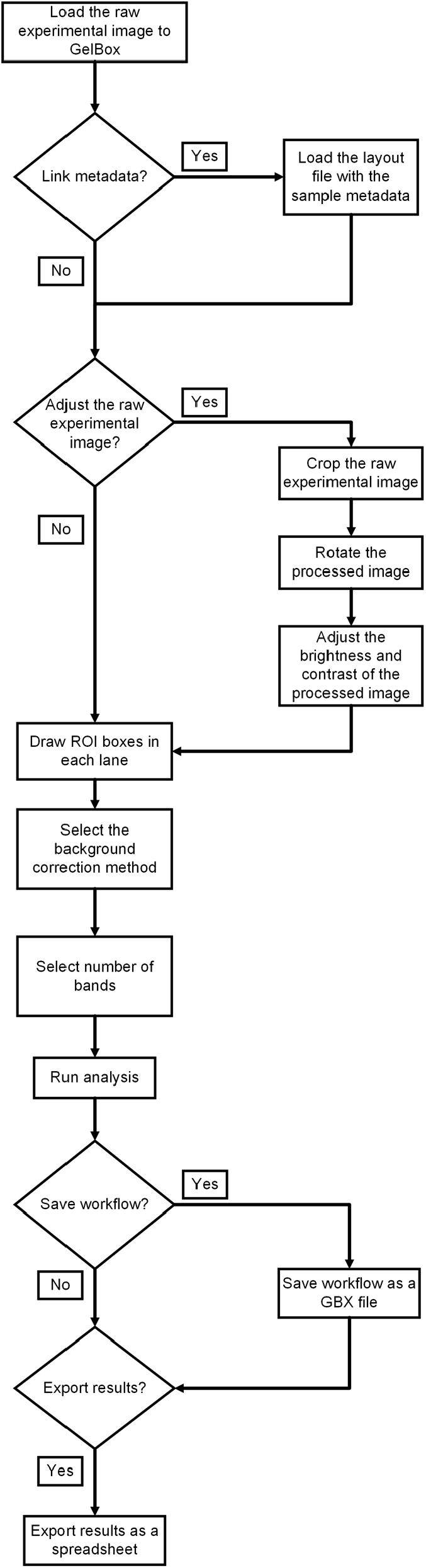
Analysis workflow.

### Metadata

Gels and immunoblots are usually associated with metadata defining each lane’s sample information, e.g., sample type, specimen number, experimental group, etc. Users can add these metadata to a GelBox analysis by uploading a spreadsheet.

### Image files and adjustments

GelBox allows users to load images stored in most commonly-used formats including TIF (Tagged Image File Format) and PNG (Portable Network Graphics). Users can crop their images to a desired size, rotate the image if necessary, and then adjust the brightness and contrast as needed. Additionally, users can invert their image if needed. GelBox does not allow users to perform gamma correction as this adjustment can change the relative intensity of bands.

### Regions of interest

Users define a region of interest (ROI) on each lane they want to analyze by drawing a rectangular box on the image. If multiple boxes are drawn, GelBox ensures they all have the same size. If the user changes the size of a box, the other boxes resize automatically. If metadata have been provided, each ROI is linked to the appropriate information.

### Analysis

Many investigators want to quantify the intensities of bands in their images. Two factors that complicate this task are background correction and potential overlap between band profiles. GelBox provides tools that handle these issues with reproducible data-driven approaches.

### Background correction

GelBox allows the user to remove trends in the background data using one of three methods: cubic smoothing spline, straight line, and rolling ball (Figure 3). The cubic smoothing spline method marks portions at the ends of the density profile and interpolates a cubic spline between these segments. The straight-line method follows a similar approach, where the sampled points are used to fit a straight line. Lastly, in the rolling ball method, a circular element is “rolled” under the density profile. The route tracked by the center of the element is used to define the background intensity (6).

**Figure 3.**
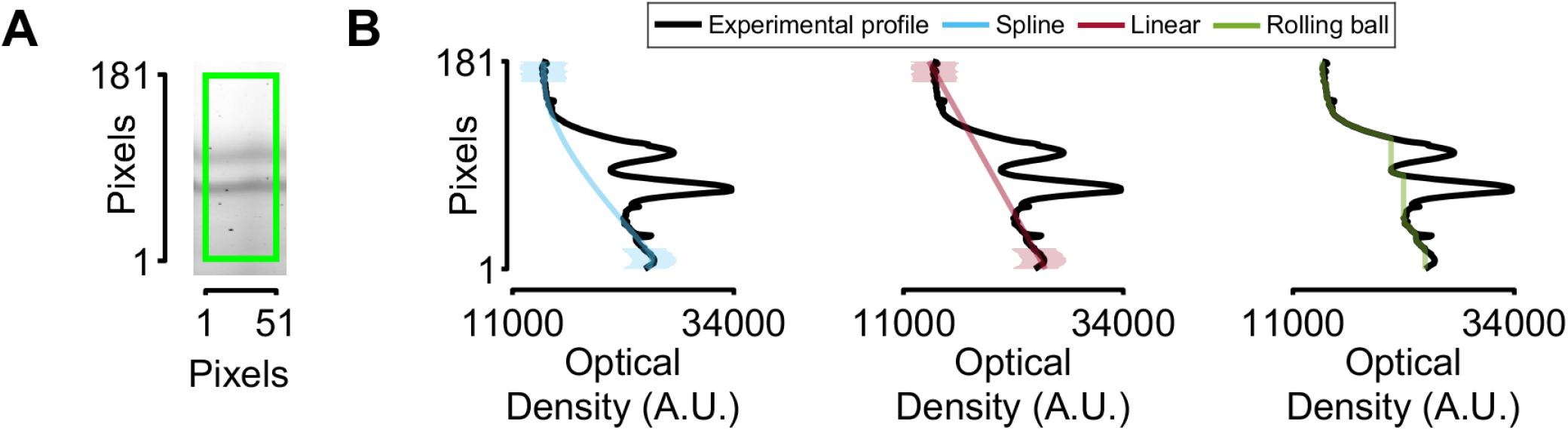
Background correction methods. (A) Region of interest from a gel showing isoforms of cardiac titin. (B) Density profile and estimated background profiles with three methods. The shaded colored areas highlight the portions at the ends of the profile that are used to estimate the background with the cubic smoothing spline and linear approaches.

### Band quantification

GelBox quantifies the total density associated with each band by fitting mathematical functions to the profile of the region of interest. Our lab has already published a variant of this method (7) and showed that it provides more accurate quantification when band profiles overlap. Specifically, GelBox fits the sum of n (potentially asymmetric) Gaussians to the band profile with a mathematical function defined as

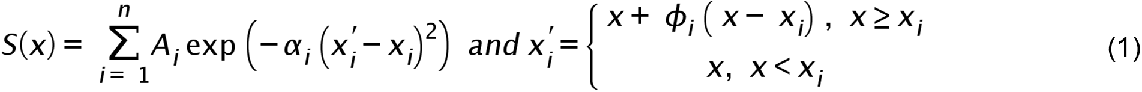

where x is the vertical position in the ROI, n is the number of bands, A_i_ and x_i_ are the peak amplitude and center of each band, α_i_ determines the width, and ϕ_i_ is a skew parameter.

The fit is performed by using the simplex search method (8) to minimize Equation 2

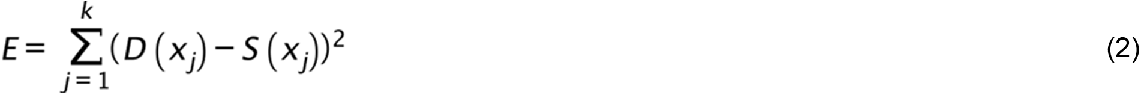

where k is the height of the ROI in pixels, and D(x) and S(x) are the density and fitted profiles, respectively. The optimization scheme allows users to impose constraints on the parameters if required. The area associated with each band is calculated by integrating the area under each Gaussian function (9).

### Workflow traceability

GelBox provides an option to combine the raw image, the user-supplied metadata, each step in the workflow, and the band data in a single GBX (short for GelBox) file. GBX files are compressed files written in MAT-format (10) and can be re-opened in GelBox to revisit analyses. This is important for rigor, reproducibility, and traceability.

### Data export

Users can also export a summary of each analysis in Excel format. The first sheet in each file combines the available metadata, the position of each region of interest, calculated densities, and the background correction method. Additional sheets provide the raw and fitted density profiles for each lane. GelBox can also generate summary figures that show the density profiles and quantification along with the raw and adjusted image.

### Availability of software

GelBox is available at no cost to academic users from https://campbell-muscle-lab.github.io/GelBox/.

The installation instructions are provided on the website. Users can access the source code through the GitHub repository (https://github.com/Campbell-Muscle-Lab/GelBox).

## RESULTS

Background correction can impact the quantification of bands on gels and immunoblots. This was investigated by analyzing computer-generated images with known band intensities superposed on mathematically-defined backgrounds. Specifically, 100 images were generated by adding a single Gaussian function to a profile defined by a third-order polynomial. The Gaussian and polynomial parameters were selected from pseudo-random distributions, yielding a wide range of qualitative image types. Speckle noise was added to each image to mimic experimental artifacts.

Each image was then analyzed using the cubic smoothing spline, linear fit, and rolling ball background correction methods (Figure 4A – B). The ratios of the calculated to actual band densities were determined (Figure 4D). One-sample t-tests showed that the linear and rolling ball correction methods led to underestimates of the true band density. Both approaches removed a portion of the true band during the correction procedure. These tests suggest that background correction via cubic smoothing splines may be more robust than linear or rolling ball methods when analyzing gels and immunoblots.

**Figure 4.**
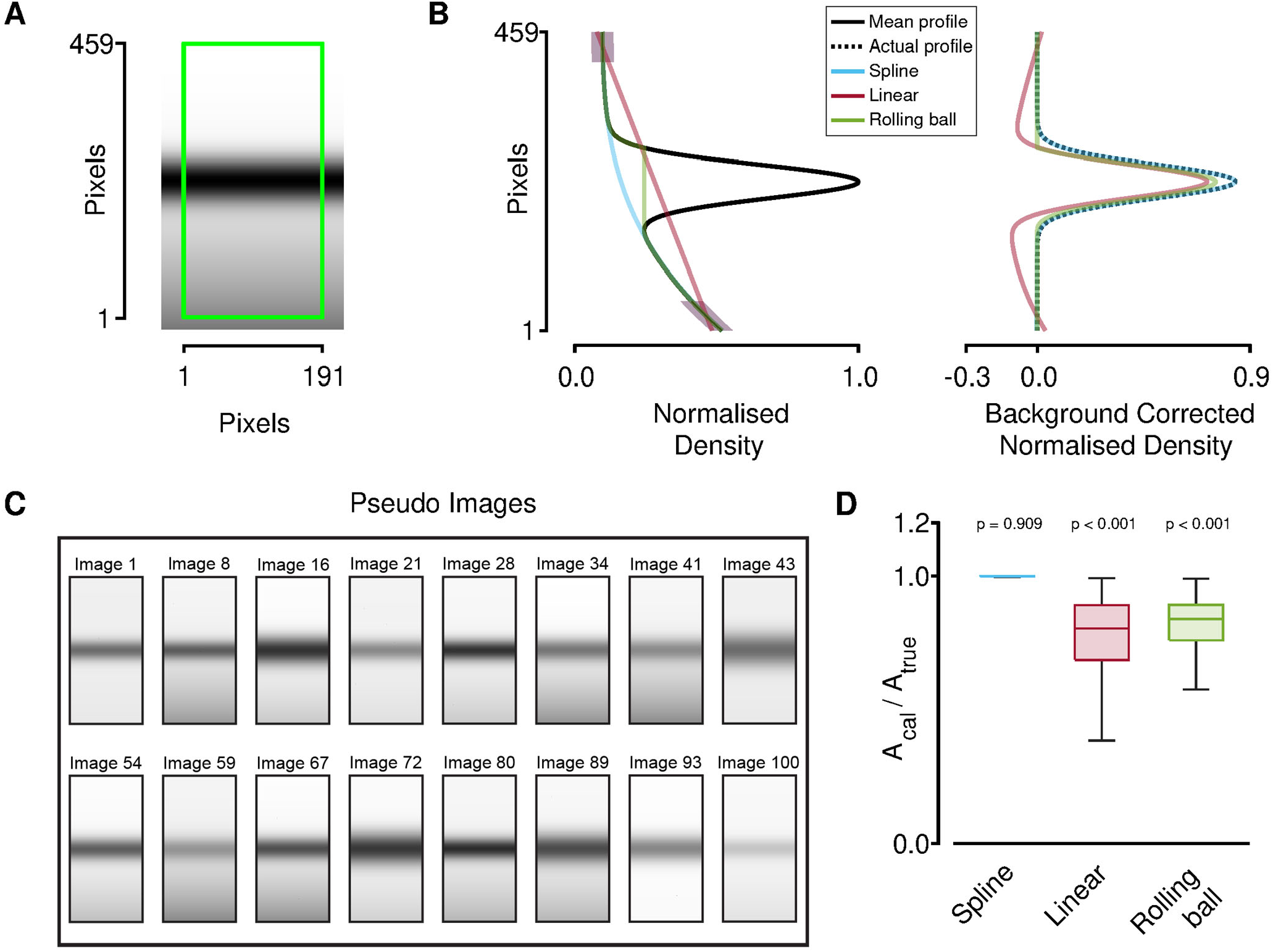
Comparison of background correction methods. A) Computer-generated pseudo image. The ROI is highlighted with the green rectangle (B) Left-hand panel: the mean density profile and the estimated backgrounds. The right-hand panel: superposed traces of corrected and the actual profile. (C) Examples of pseudo images. (D) The ratios of the calculated areas to actual areas from 100 pseudo images. p-values show the results of one-sample t-tests that compare the ratios calculated by each method to unity. The linear and rolling ball methods significantly underestimated the true intensity of the simulated bands. Plots show extremes (end-caps), quartlles (box), and median (central bar).

## DISCUSSION

GelBox is a computer program tailored to help researchers analyze gels and immunoblots. It allows the user to complete their analysis workflow in a single package and saves the raw data, user-defined metadata, and all workflow steps in a single file. This can enhance rigor, reproducibility, and traceability.

Another strength is the use of curve-fitting to determine band densities. As previously explained (7), this is particularly advantageous when the band profiles overlap. The fitting procedures reduce unintended bias and add experimental rigor. Three different background correction methods are provided. The linear and rolling ball approaches are implemented in many other packages but our mathematical analysis suggests that they may lead to underestimates of band intensities (Fig 4). Cubic smoothing splines may provide a better alternative.

A limitation of GelBox is that it is written in the MATLAB language. Although the code is open source, MATLAB is a commercial product. Many universities have site licenses, so academics can use and potentially adapt the software for their own use without financial cost. Users can also install a standalone pre-compiled version of GelBox. This would allow them to use GelBox without installing the main MATLAB package. However, this approach requires installing MATLAB runtime prerequisites.

Although free, these are cumbersome to install and complicated to maintain.

The website for the software provides comprehensive tutorials that explain how to analyze images with GelBox. The site also provides a troubleshooting guide that describes how to mitigate commonly encountered issues.

GelBox simplifies the analysis of gel and immunoblot data, but it cannot compensate for a poorly conducted experiment. The quality of the raw data is critical. In addition, users need to have a clear understanding of how GelBox works to analyze their data appropriately.

## ACKNOWLEDGMENTS

The authors would like to thank Nicholas S. LaFave (University of Massachusetts, Lowell) and Peter Omondi Awinda (Washington State University) for providing feedback on an early version of GelBox.

This work was supported by NIH grants HL149164 and HL148785.

